# Chemosensory structure and function in the filarial nematode, *Brugia malayi*

**DOI:** 10.1101/427229

**Authors:** Lisa M. Fraser, R. Isai Madriz, Divyaa Srinivasan, Mostafa Zamanian, Lyric C. Bartholomay, Michael J. Kimber

## Abstract

Nematode chemosensory behaviors underlie fundamental processes and activities in development, reproduction, tropisms and taxes. For parasitic species, chemosensation is essential for host seeking and host and tissue invasion behaviors. Such fundamental biology presents an attractive target for developing behavior-blocking anthelminthic drugs, but the anatomy and functional relevance of parasitic nematode chemosensory machinery are poorly understood. The goals of this study were to better understand the chemosensory apparatus and behaviors of infectious stage *Brugia malayi* (Spirurida: Onchocercidae), a mosquito-borne nematode and etiological agent of Lymphatic Filariasis. Scanning electron microscopy revealed that amphids, the major chemosensory organs, are present on adult *B. malayi* and arranged in a conserved manner. Internal sensory neuroanatomy display structural differences between life stages, and a simpler chemosensory architecture as compared to free-living nematodes. Positive and negative chemotactic behaviors were identified for a repertoire of chemicals with known chemostimulatory activity for the mosquito host that may facilitate host-selectivity and invasion. This is the first description of chemosensory anatomy and chemotactic behaviors in *B. malayi* that reveal the involvement of chemosensation in parasite transmission and host invasion.

Key findings
• Amphidial arrangement in *B. malayi* is less complex than free-living nematodes.
• Chemosensory neuroanatomy is stage-specific and simpler than free-living nematodes.
• *B. malayi* responses to stimuli can be measured using a plate-based assay.
• Chemostimulants associated with mosquito host-seeking induce negative and positive tropisms for L3 stage *B. malayi*.

## Introduction

Chemosensation is an essential behavior used by multi-cellular organisms to receive and respond to chemical cues in the external environment (Bargmann, 2006b; Prasad and Reed, 1999). This process begins with the detection of environmental signaling molecules by chemoreceptors, most commonly G protein-coupled receptors (GPCRs) (Prasad and Reed, 1999). Receptor binding leads to the activation of chemosensory neurons and downstream motor neurons, resulting in a physical response to the stimulus (Bargmann, 2006b). Although this fundamental schema for chemosensation is generally conserved across organisms, the chemosensory systems of insects, nematodes and all higher animals reveal significant adaptive radiation in the chemosensory machinery. In *Caenorhabditis elegans* adult hermaphrodites, 32 neurons are presumed to be chemosensory neurons (Thomas and Robertson, 2008; Troemel *et al*., 1995; Ward *et al*., 1975), as compared to 2600 olfactory neurons in *Drosophila melanogaster*, and mammals that have millions of odorant sensory neurons (Bargmann, 2006b; Liberles, 2014; Li and Liberles, 2015). Despite having a relatively simple chemosensory neuronal architecture, the chemoreceptor complement in *C. elegans* is complex such that more than 1000 putative chemoreceptors are encoded in the genome (Bargmann, 2006b; Robertson and Thomas, 2006). A facile summary could be that nematode chemosensory systems are anatomically simple but molecularly complex, while those of insects and higher animals are anatomically complex but molecularly simple. The expression of many chemoreceptors per chemosensory neuron in nematodes provides the organism with an efficient mechanism to detect a wide array of compounds with remarkable precision within the confines of a simple neuronal architecture (Bargmann, 2006b; von der Weid *et al*., 2015).

Nematode chemosensory behavior has been characterized chiefly through the study of *C. elegans*, a free-living model species (Bargmann and Horvitz, 1991; Troemel *et al*., 1995; Troemel *et al*., 1997; Chalasani *et al*., 2007). Olfactory cues drive a number of important behaviors in *C. elegans*, including food and mate seeking behavior, avoidance of noxious conditions and entry/exit into the dauer stage (Troemel *et al*., 1997; Bargmann, 2006b). The chemosensory capacity and behavior of parasitic nematodes and is much less well understood, but it is clear that chemosensation is critical for host seeking and host invasion behaviors (Hallem *et al*., 2011; Chaisson and Hallem, 2012; Castelletto *et al*., 2014). The infective stages of skin penetrating parasitic nematodes such as *Strongyloides ratti* and *Ancylostoma caninum* exhibit positive chemotaxis to host serum, which may provide a positive stimulus for skin penetration and thereby facilitate transmission and infection (Vetter *et al*., 1985; Granzer and Haas, 1991; Koga and Tada, 2000). In addition, studies in both plant- and animal-parasitic nematodes have shown that chemosensation underlies host seeking and host preference (Robinson and Perry, 2006; Hallem *et al*., 2011; Wang *et al*., 2011; Castelletto *et al*., 2014). Collectively, these findings support the hypothesis that the capacity to perceive and respond to stimuli is important for the transmission of, and establishment of infection by, parasitic nematodes.

Lymphatic Filariasis (LF) is a disease caused by mosquito-borne filarial nematodes including *Wuchereria bancrofti* and *Brugia malayi*. An estimated 120 million people are infected and despite an orchestrated and widespread mass drug administration (MDA) program, LF remains a significant global health concern (WHO 2017). The MDA strategy relies on anthelmintics that reduce the burden of juvenile stage parasites in the infected host in order to prevent transmission to mosquito vector species, i.e., MDA is not curative for adult-stage parasites. As such, there is a real and urgent need to develop new chemotherapeutics with new modes of action and efficacy against additional life stages to prevent establishment of infection or kill adult-stage parasites. For successful infection, infective L3 stage filarial parasites, which are deposited on the skin while the vector mosquito is taking a blood meal, must identify and penetrate the human host via the wound made by the mosquito mouthparts (Bartholomay, 2014). The parasites then migrate to the lymphatic vasculature where they undergo an additional two molts to become sexually mature adults. Accurate perception of host-derived chemical stimuli likely facilitates this complex invasion process, contributing to host selectivity and migration to appropriate end tissues. Disruption of chemosensory perception, therefore, may prevent filarial parasites establishing infection and developing to the adult stage, which is generally recalcitrant to anthelmintics. Exploitation of this paradigm requires a better understanding of chemosensory behavior in filarial nematodes. To this end, research has been limited to the infectious L3 stage of *B. pahangi*, which is attracted to host serum and sodium ions in a concentration dependent manner (Gunawardena *et al*., 2003; Kusaba *et al*., 2008; Mitsui *et al*., 2012). The goals of the present study were to characterize the ultrastructure of chemosensory apparatus of human-infecting filarial worm parasites and to profile functional chemosensory behavior in the invasive L3 stage.

## Materials and methods

### Mosquito maintenance

Adult female *Aedes aegypti* (Black eyed Liverpool strain, LVP) were used for all experiments. This species is not a natural vector for *B. malayi*, but this strain was selected for susceptibility and is the model system for studies of filarial worm-mosquito interactions (MacDonald, 1962). Adult mosquitoes were maintained on a diet of 10% sucrose and in an environmental chamber with controlled conditions (27°C ± 1°C and 75% ± 5% relative humidity) and a 16:8 photoperiod.

#### *Establishment of* Brugia malayi *infection and parasite collection*

Cat blood containing *B. malayi* microfilariae (mf) was obtained from the University of Georgia NIH/NIAID Filariasis Research Reagent Resource Center (FR3) and diluted with defibrinated sheep blood (Hemostat Laboratories, CA, USA) to achieve a concentration of 150–250 mf per 20 μl. To establish infection, three- to five-day old female *Ae. aegypti* (LVP) were allowed to feed through a Parafilm (Bemis NA, Neenah, WI, USA) membrane on blood held in a water-jacketed glass membrane feeder with circulating water heated to 37°C. Mosquitoes were sucrose-starved for 24 hrs prior to blood feeding and those that did not take a blood meal were removed. Infected mosquitoes were maintained under controlled conditions for 12–19 days post infection (dpi) to allow development of parasites to the L3 stage. To collect L3 stage parasites, infected adult female *Ae. aegypti* were cold-anesthetized on ice and dissected into three sections (head/proboscis, thorax and abdomen) in separate drops of room temperature *Aedes* physiological saline (Hayes, 1953); 154 mM NaCl, 1.36 mM CaCl_2_, 2.68 mM KCl, 1.19 mM NaHCO_3_, and parasites were allowed to migrate out of the mosquito into the saline solution. In addition to L3 parasites harvested in-house, we also obtained supplementary L3, L4, adult male and adult female stages from FR3 for scanning electron microscopy, staining and chemotaxis plate assays. For these purposes, no difference was observed between L3s freshly collected from infected mosquitoes and those obtained from FR3.

#### Scanning electron microscopy

Parasites were fixed in 1.5 mL microcentrifuge tubes with 2% paraformaldehyde and 2% glutaraldehyde (Sigma-Aldrich, St. Louis, MO, USA) in 0.1 M cacodylate buffer (Thermo Fisher Scientific, Waltham, MA, USA), pH 7.2, for 24 hrs at 4°C. Specimens were rinsed four times with the same buffer and post-fixed in 2% aqueous osmium tetroxide (Sigma-Aldrich) for 24 hrs before being rinsed four times with the same buffer followed by dehydration in a graded ethanol series (10, 30, 50, 70, 90, 95, 100, 100, 100%; each step for 15 min). Finally, samples were dried using ultrapure ethanol and carbon dioxide in a critical point drying apparatus (Denton Vacuum DCP-1, Denton Vacuum, LLC, Moorestown, NJ USA). Dried samples were stored in a desiccator or adhered to aluminum stubs using double-stick tape and silver paint. The samples were sputter-coated (Denton Desk II sputter coater, Denton Vacuum) with 120 nm palladium-gold (60:40). Samples were viewed with a scanning electron microscope (JEOL 5800LV) at 10–13 kV in the Microscopy and NanoImaging Facility, Iowa State University. Images were captured using a SIS ADDA II (Olympus Soft Imaging Systems Inc., Lakewood, CO, USA).

#### Dye filling assays

Staining of live parasites (L3, L4, adult male and adult female) was performed using 1,1’-dioctadecyl-3,3,3’,3’-tetramethylindocarbocyanine perchlorate (DiI, Thermo Fisher Scientific, Waltham, MA, USA). Worms were washed using RPMI 1640 (Thermo Fisher Scientific) containing pen-strep (0.4 units penicillin/ml, 0.4 mcg streptomycin/ml). *B. malayi* were then placed into clean RPMI 1640 containing DiI (10 μg/ml) and pen-strep and incubated at 37°C for 16–24 hrs. After staining, animals were washed twice with RPMI 1640 containing pen-strep and visualized using a Nikon Eclipse 50i compound fluorescent microscope (Nikon, Japan).

#### Chemotaxis plate assays

Chemotaxis assays developed for *Caenorhabditis elegans* (Margie *et al*., 2013) were adapted and refined for use with *B. malayi* L3. Briefly, sterile 35 mm petri dishes (hereafter referred to as "plates”) containing 0.8% agarose (Fisher Bioreagents). Plates were divided into four equal quadrants. A 3 mm diameter circle was marked around the point of intersection of the perpendicular lines to designate the center. In each quadrant, an additional circle, 3 mm in diameter, was drawn 5 mm from the plate center. Circles in diagonally opposite quadrants were marked as Test 1 (T1) and Test 2 (T2), and the other two circles were marked Control 1 (C1) and Control 2 (C2) (Fig. 1). The chemotaxis assay was developed using 3 μl of each undiluted test compound (except 3-methyl-1-butanol) placed in circles T1 and T2, and 3 μl of 1xPBS placed in circles C1 and C2. The exception was 3-methyl-1-butanol, which was diluted in mineral oil in a 1:1 ratio; for this compound, 6 μl of solution was placed in circles T1 and T2 and 3 μl of mineral oil was placed in the control circles. Two sets of control plates were also prepared. For one set of control plates, 3 μ! of 1xPBS was placed in all four circles (Test and Control), and for the other set 3 μl of mineral oil was placed in all four circles. Once all the compounds were pipetted out as mentioned above, the plates were allowed to sit at room temperature for 1 hour. For each test and control replicate, eight L3s were placed in 5 μl 1xPBS at the origin of the plate. Once the L3s were placed at the origin, the plates were incubated in a humidified chamber at 35°C for at least 30 min.

**Fig. 1.**
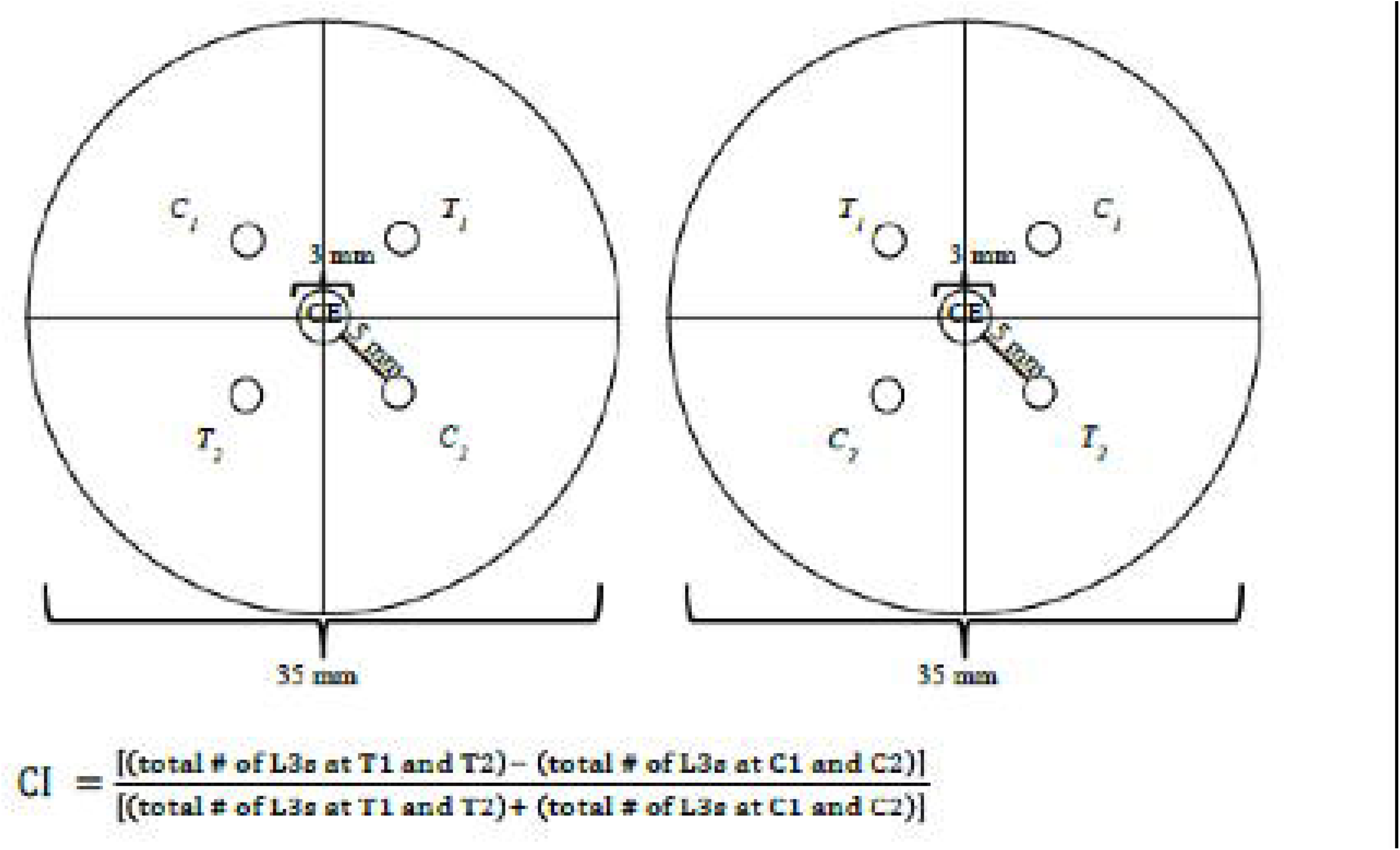
*B. malayi* chemosensory behaviors are profiled using plate-based chemotaxis assays. Schematic of chemotaxis assay used. Infectious L3 *B. malayi* are placed in the center (CE) and allowed to distribute over the plate. After 30 minutes the number of parasites in T1, T2 (test compounds), C1 and C2 (control compounds) are counted and the chemotaxis index (C.I.) is calculated as indicated. Two plate arrangements were used to minimize any external or directional bias.

After incubation, L3s in each quadrant were counted. Only the L3s in a quadrant received a score and those L3s that did not leave the center or were on the line between two quadrants, did not receive a score. Three trials were performed for each compound tested, with an average of 20 plates per trial per condition. Using these data, a chemotaxis index (CI) for each test and control plate was calculated as:

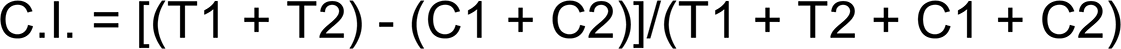

Scoring for maximum repulsion and maximum attraction ranged between -1.0 and +1.0, respectively. Plates where at least four parasites moved from the center were retained for calculation of CI. This was used to filter plates where the majority of worms remained trapped together in a pocket of liquid at the center. Compounds that elicited strong attraction or repulsion were re-tested in a different lab environment, controlling for differences in mosquito colony, environmental parameters, and diluent. Five independent trials were carried out with an average of three plates per trial per condition.

All liquid compounds were tested in an undiluted state, solid compounds were prepared at a concentration of 1 M. Compounds used in these assays included: Human AB Serum, (MP Biomedicals), NaCl (Fisher Bioreagents), L-Lactic acid (undiluted)(Sigma-Aldrich), 3-methyl-1-butanol (Sigma-Aldrich), 1-nonanol (Sigma-Aldrich), Fetal Bovine serum (Fisher Scientific), Mineral oil, and PBS (DOT Scientific Inc).

#### Statistical analysis

For analysis of chemotaxis plate assays, unpaired two-tailed t-tests were used to compare CI of test compounds to CI of control compounds. Each chemotaxis plate was considered a replicate with worms originating from a mixed population of mosquitoes, with additional replication through staggered mosquito infection trials. No significant batch effects were observed across staggered trials. All statistical analyses and visualization were carried out using the statistical programming language R.

## Results

### B. malayi *external chemosensory structures*

The primary chemosensory organs in nematodes are the amphids, a pair of lateral sensory structures located anteriorly with cephalic or cervical external openings either side of the mouth of the nematode (Ward *et al*., 1975; Wright, 1983; Perkins *et al*., 1986; Robinson and Perry, 2006; Bacaj *et al*., 2008). Amphid pore shape is highly variable among nematodes and while the significance of this is unknown, it may be that the more convoluted shapes are functional and enhance the sensitivity of these organs as chemoreceptors. Amphidial structure in filarial nematodes is poorly defined so we used scanning electron microscopy (SEM) to describe the external arrangement of amphids across the *B. malayi* life cycle. SEM confirmed the presence of amphids with an arrangement conserved in all life stages examined (L3, L4, adult male and adult female) (Fig. 2). *B. malayi* amphid pore structure is crescent-shaped and generally appeared less prominent than those found in *C. elegans* but similar to what has been observed in both *Onchocerca lupi* and *O. eberhardi*, two species more closely related to *B. malayi* (Uni *et al*., 2007; Mutafchiev *et al*., 2013). Although not the focus of this study, the arrangement of other anterior sensillae was also noteworthy. *B. malayi* possess fewer cephalic papillae (four inner labial sensilla and four outer labial sensilla) (Fig. 2) than *C. elegans* (six inner labial sensilla, six outer labial sensilla and four additional cephalic sensilla), which is considered to have the classical arrangement of cephalic papillae (Ward *et al*., 1975; Tippawangkosol *et al*., 2004). Like *B. malayi, O. eberhardi* exhibits a reduction in anterior papillae (Uni *et al*., 2007) suggesting that this reduction may be a common feature of filarial worms.

**Fig. 2.**
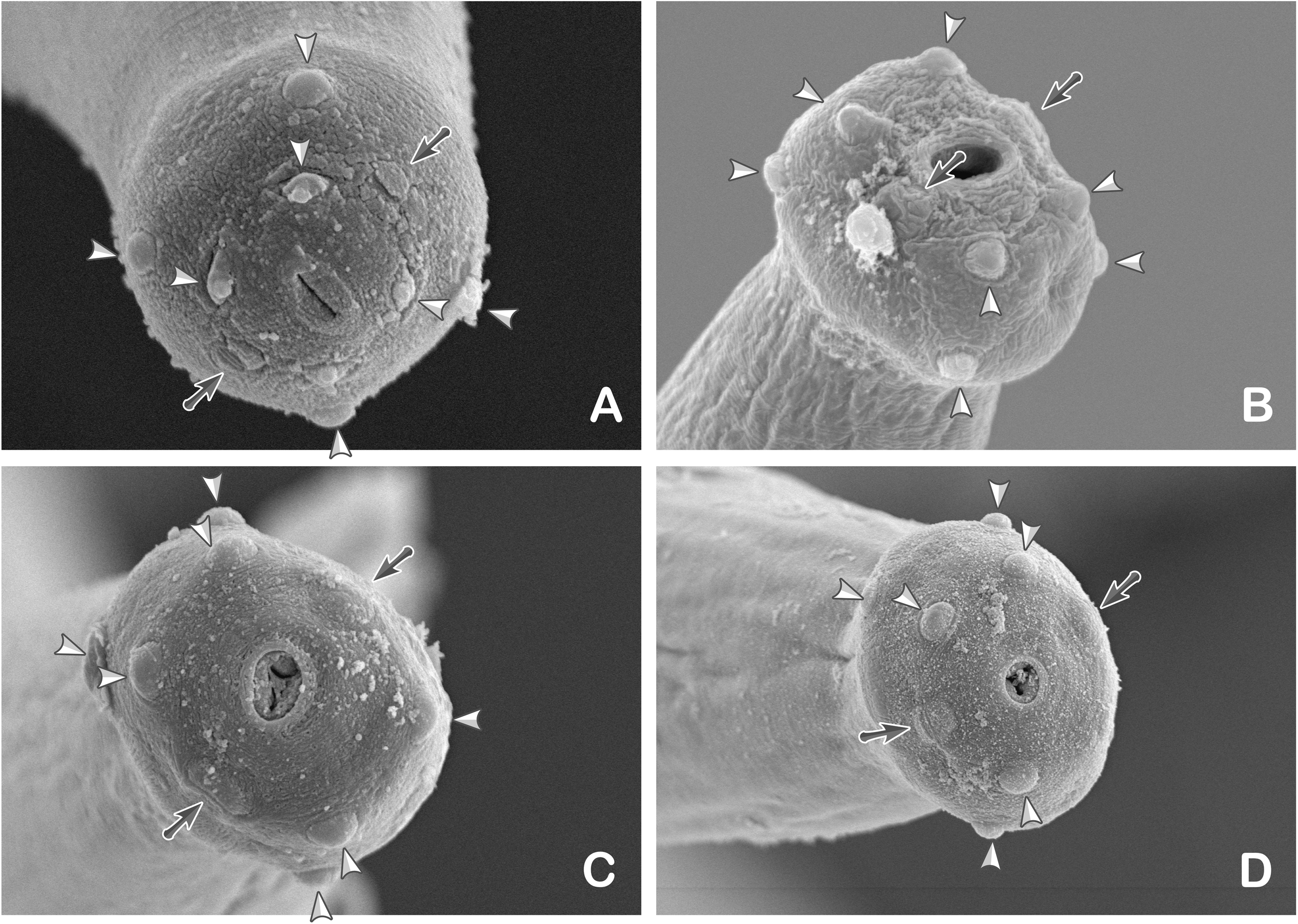
*B. malayi* parasites have conserved sensory ultrastructures across multiple life stages. Scanning electron images of the anterior portion of (A) L3, (B) L4, (C) adult male and (D) adult female *B. malayi* are shown. These data reveal that all life stages examined possess amphid pores (arrows) and cephalic papillae (arrow heads). Scale bars: 5 μm (A and B), 10 μm (C and D).

### B. malayi *chemosensory neuroanatomy displays stage-specific variation*

To characterize *B. malayi* sensory neuroanatomy, we used a facile dye-filling assay utilizing the lipophilic dye 1,1’-dioctadecyl-3,3,3’,3’-tetramethylindocarbocyanine perchlorate (DiI) that results in fluorescent loading of the sensory neurons exposed to the solution (Hedgecock *et al*., 1985). When applied to *B. malayi*, we observed dye-filling of anterior neurons throughout all life stages examined (L3, L4, adult male and female), however, dye-filling patterns varied according to life stage. The amphid channel, generated by two glia cells called socket and sheath cells and filled with a gelatinous matrix exposed to the environment (Ward *et al*., 1975; Wright, 1983), consistently stained brightly in all life stages and could be readily discerned (Fig. 3). Sheath glia cell bodies were clearly visible in L3, adult male and female preparations but not in the L4 stage parasites (Fig. 3B, H, K). These cells line the posterior amphid channel and are required for the proper function of amphid sensory neurons (Bacaj *et al*., 2008). Socket cell bodies were consistently visualized in L3 stage parasites (Fig. 3B) but not in other stages. Socket cells line the anterior portion of the amphid and surround the terminal tips of ciliated sensory neurons. Finally, retrograde loading of DiI progressed to the circumpharyngeal nerve ring in all stages except adult males (Fig. 3B, E, K). Noticeably, no amphidial neurons were observed to take up DiI in any *B. malayi* life stage (Fig. 3). This is a distinct departure from *C. elegans* where six amphid neurons (ASH, ASI, ASJ, ASK, ADL and AWB) are visible using this technique (Srinivasan *et al*., 2008). Although the neuroanatomy of nematodes is considered highly conserved in general, these differences in dye-filling patterns of chemosensory neuroanatomy may indicate variation in the neuroanatomy of this functional system.

**Fig. 3.**
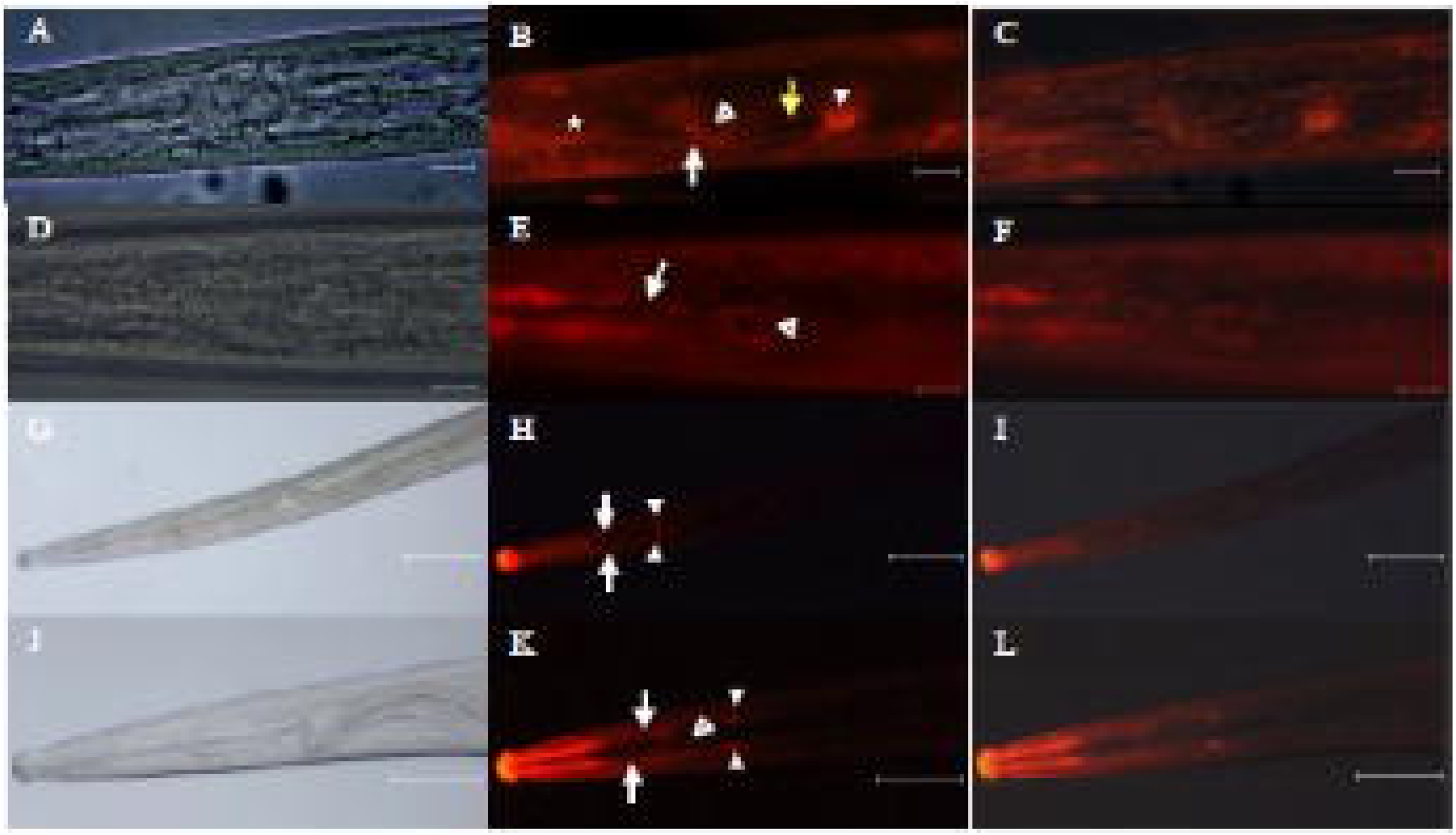
DiI staining reveals sensory neuroanatomy of *B. malayi*. Fluorescent images of anterior sensory structures in *B. malayi* using DiI stain show sheath cell bodies (closed arrowheads), nerve ring (open arrowheads), dendrites (white arrows), axons (yellow arrow) and socket cell bodies (star). Images are from L3 (A-C), L4 (D-F), adult male (G-I) and adult female (J-L) *B. malayi*. A-F show lateral views while G-L show dorsoventral views. Total magnification: 900x (A-F) and 150x (G-L). Scale bars: 10 μm (A-F) and 100 μm (G-L).

### B. malayi *chemosensory responses can be interrogated using a plate-based assay revealing attraction to simple and complex gustatory stimuli*

A number of neurons are involved in nematode chemosensory responses such that each type of neuron recognizes different categories of molecules (Smith *et al*., 2013). Gustatory cues including salts, amino acids and other water-soluble molecules are recognized by the ASE amphid neurons, the primary taste receptor neurons in *C. elegans* (Bargmann and Horvitz, 1991; Bargmann, 2006a; Smith *et al*., 2013). Sodium chloride has been shown to elicit behavioral responses in *C. elegans* demonstrating that this is a relevant gustatory compound perceived by nematodes (Ward, 1973). We hypothesized that *B. malayi* would also be able to perceive environmental NaCl and, like *C. elegans*, would exhibit motile behaviors in response. To test this hypothesis we refined a agar plate-based assay (Fig. 1) to measure *B. malayi* responses to a range of applied stimuli, using NaCl for both assay development and as a model gustatory compound. Using our assay, we found *B. malayi* L3s were strongly attracted to 1M sodium chloride (NaCl) (mean C.I. = 0.55, p < 0.01) when compared to water controls (Fig. 4). It should be noted that although the concentration of NaCl applied to the plate was high (1M), the assay allows for the diffusion of water-soluble compounds through the agar, thereby creating a concentration gradient radiating from the application point. It is likely that parasites were moving to an "ideal” concentration within that gradient. That said, previous studies have shown that when placed in a low NaCl concentrations, the skin-penetrating parasite *Stmngyloides stercoralis* will migrate up a NaCl gradient to concentrations as high as 1.1M (Forbes *et al*., 2004).

Having established a straightforward assay that allows us to describe *B. malayi* responses to simple gustatory stimuli, we wanted to further profile filarial chemosensory behavior, focusing on the role chemosensation plays in parasite invasion and disease transmission. Specifically, we wanted to examine filarial responses to complex host derived biofluids, or solutions recapitulating such fluids, of which NaCl is a component. Attraction to such biofluids may drive host invasion. *Aedes* saline is a physiological saline containing 154 mM NaCl that is designed isotonic to mosquito hemolymph (Hayes, 1953). *Brugia* L3 were strongly attracted to *Aedes* saline (mean C.I. = 0.28, p < 0.05) (Fig. 4). Although more complex than a simple NaCl solution, this positive tactic response to *Aedes* saline may be a function of the same NaCl response previously observed and there was no significant difference between the two responses. To test parasite behavior to true host biofluids, we used commercially available human and fetal bovine sera. *Brugia* L3 were strongly attracted to both the human serum (mean C.I. = 0.46, p < 0.01) and the fetal bovine serum (mean C.I. = 0.59, p < 0.01) (Fig. 4). Although the positive response was stronger than that observed to *Aedes* saline, there was no statistically significant difference between the three responses. It has been suggested that differential parasite responses to chemical cues may reflect host-specificity (Vetter *et al*., 1985; Granzer and Haas, 1991; Koga and Tada, 2000; Kusaba *et al*., 2008; Castelletto *et al*., 2014), however, the quantitatively similar attractive responses of *B. malayi* L3 to bovine and human serum suggest any host discriminating sensory cues are not present in these fluids as bovids are not hosts for *B. malayi*.

**Fig. 4.**
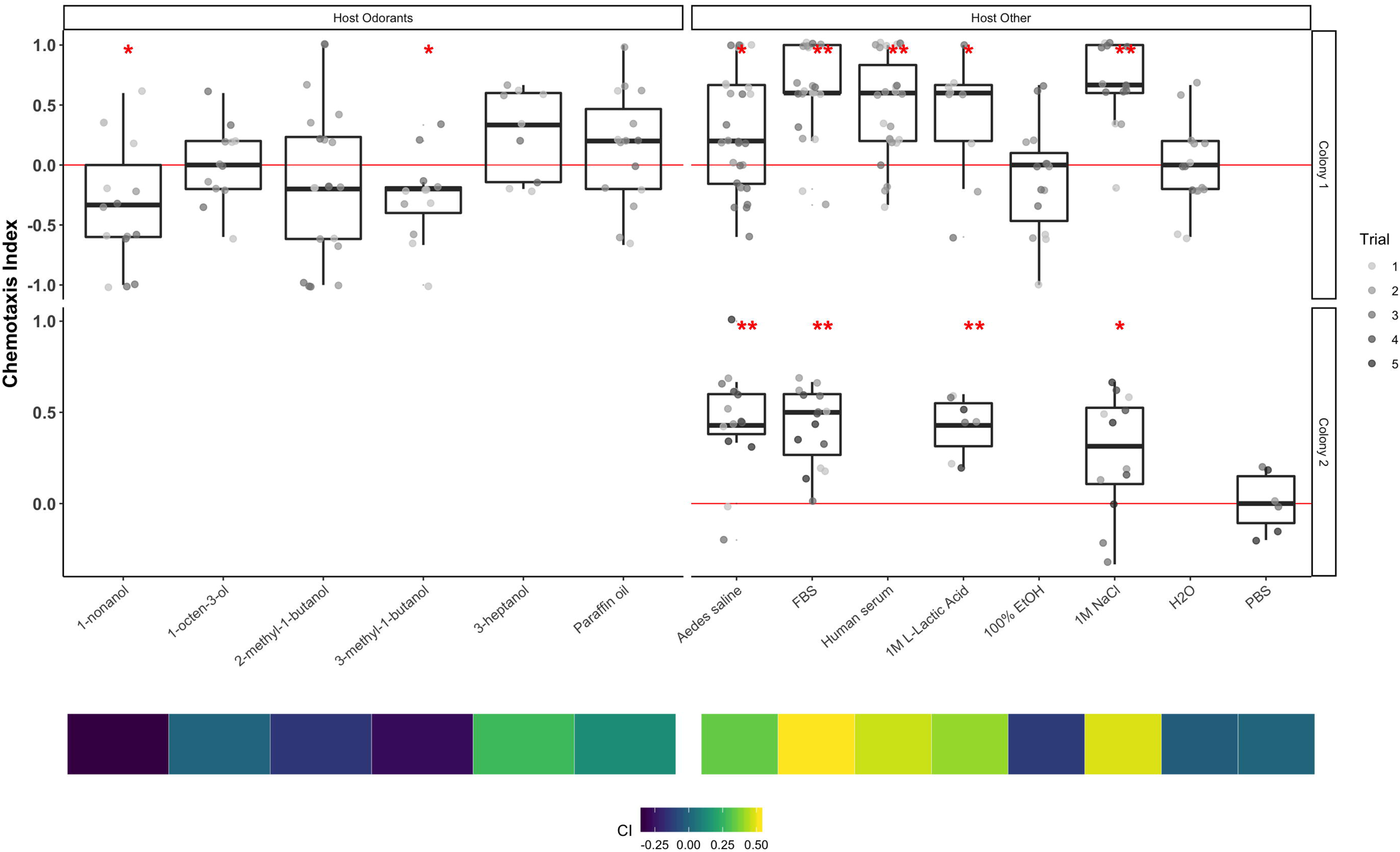
*B. malayi* parasites exhibit robust chemoattraction and chemorepulsion behaviors in response to host-associated compounds. *B. malayi* L3 stage worms are attracted to *Aedes* saline, mammalian serum (FBS and human serum), NaCl, and L-lactic acid. Infective stage larvae show aversion to 1-nonanol and1-octen-3-ol. Control diluent for compounds in the left and right panels are parrafin oil and H_2_O, respectively. Box-plots are shown with significance relative to control (two-tailed t-test: ** p-value < 0.01; * p-value < 0.05). Each data point represents a chemotaxis plate. Three independent trials were carried out with an average of 20 plates per trial per condition in the first set of experiments. The most significant chemotactic responses are recapitulated using parasites produced from a different mosquito colony in a different laboratory environment ("Colony 2”), and with an alternative diluent (PBS instead of H_2_O). Each data point represents a chemotaxis plate. Five independent trials were carried out with an average of three plates per trial per condition. Box-plots are shown with significance relative to control (two-tailed t-test: ** p-value < 0.01; * p-value < 0.05). A heatmap (bottom) shows mean C.I. across all experiments for each compound tested.

#### Host-derived olfactory compounds elicit specific tactic behaviors in L3 stage B. malayi

Like gustatory compounds, odorants (volatile compounds) elicit behavioral responses in *C. elegans* but these responses are mediated by a different subset of chemosensory neurons, the olfactory neurons AWA, AWB and AWC (Bargmann, 2006a; Chalasani *et al*., 2007). Studies using skin-penetrating parasitic nematodes including S. *stercoralis* suggest that perception of odorants plays an important role in host-selection and invasion, likely more so than perception of gustatory cues (Castelletto *et al*., 2014). Our hypothesis was that odorant compounds stimulate motile behaviors in infective stage *B. malayi* and contribute to host invasion in this medically important species. To test this hypothesis we used the chemotaxis plate assay to investigate L3 stage parasite responses to a select panel of host-derived odorant compounds. L-lactic acid is an odorant generated in human muscle and which is generally repellant to free-living nematodes and neutral to mildly attractive to those parasitic species examined (Castelletto *et al*., 2014). *B. malayi* L3 were attracted to L-lactic acid (mean C.I. = 0.42, p < 0.05) (Fig. 4); given the caveat of variability between assay platforms, this attraction was stronger than has previously been reported for parasitic nematodes. Ethanol is an odorant that elicits broad responses in nematodes, ranging from mildly repellant *(S. stercoralis)* to strongly attractive *(C. elegans)*. Here, ethanol was mildly repellant to *B. malayi* L3 but this response was not significant (mean C.I. = -0.12, p = 0.12). Next, we focused on select odorant components of human skin emissions. *B. malayi* L3 were repelled by 3-methyl-1-butanol (mean C.I. = -0.28, p < 0.05) (Figure 4) and 1-nonanol (mean C.I. = -0.33, p < 0.05). Although mildly repelled by 2-methyl-1-butanol, this response was not significant (p = 0.15). Responses to 3-heptanol were mildly attractive but not significant compared to vehicle (mean C.I. = 0.27, p = 0.43).

## Discussion

This study describes chemosensory apparatus and behavior in the filarial nematode, *B. malayi*, demonstrating that these parasites possess the major structures required for sensory perception and exhibit specific tactic behaviors when exposed to different chemical compounds. Amphids are the major anterior sensory organs of nematodes. The pore shape of the amphids is highly variable among nematodes, ranging from simple slits to large complex structures (Ashton *et al*., 1999). We used scanning electron microscopy to reveal all *B. malayi* life stages examined (L3, L4, adult male and female) possess simple, crescent shaped amphidial openings that are similar to amphid pores of a closely related species, *Onchocerca eberhardi* and less prominent than those of *C. elegans* (Bargmann and Horvitz, 1991). The simplicity of amphid pore shape in *B. malayi* could be a functional adaptation to the niche this nematode occupies. *C. elegans* has a very sensitive chemosensory response and is capable of detecting and responding to hundreds of different chemical cues (Troemel *et al*., 1997). In fact, *C. elegans* chemosensation is so sensitive that these nematodes are able to distinguish between bacteria that are food sources and those that are pathogenic (Pradel *et al*., 2007). Furthermore, these worms also respond to environmental compounds in a concentration-dependent manner and exhibit adaptive behavior when pre-exposed to chemical compounds (Jansen *et al*., 2002; Matsuki *et al*., 2006). Such as complex chemosensory capability may require complex anatomical features. By contrast, *B. malayi* is a parasitic nematode that infects humans and is vectored by mosquitoes so there is effectively "free-living” stage that requires an ability to discern such a multiplicity of environmental stimuli. As a result, *B. malayi* is unlikely to require specific anatomical adaptations that would enhance the acuity of the chemosensory system in order to thrive in its ecological niche. The amphids of animal-parasitic nematodes (APNs) are often less pronounced than those of free-living nematodes, thus providing support for this hypothesis (Goater *et al*., 2013).

We used the fluorescent lipophilic dye DiI (1,1’-dioctadecyl-3,3,3’,3’-tetramethylindocarbocyanine perchlorate) to visualize internal sensory neuroanatomy across multiple life stages. Our results showed stage-specific variations in dye-filling patterns, for example, while amphid channels were found to dye-fill in all life stages examined (L3, L4, adult male and female), dye-filling of amphid associated socket cell bodies was only observed in L3 stage worms. Other stage-specific variations and comparative differences to dye-filling patterns in *C. elegans* were noted (Chalasani *et al*., 2007). Traditionally it was thought that the neuroanatomy of nematodes in the class Chromadorea (which includes *C. elegans* and *B. malayi)* is highly conserved, so the variation in dye-filling patterns was perhaps unexpected although not unprecedented (Granzer and Haas, 1991; Troemel *et al*., 1997; Bargmann, 2006a; Chalasani *et al*., 2007; Hallem *et al*., 2011; Chaisson and Hallem, 2012; Castelletto *et al*., 2014). Previous studies using DiI have shown that parasitic nematodes exhibit variation in dye-filling patterns (Vetter *et al*., 1985; Koga and Tada, 2000), for example, only a single pair of amphid neurons dye-filled in the L1 stage of the APN *Parastrongyloides trichosuri* (Vetter *et al*., 1985). More recently, Han **et al**. (2015) compared dye-filling patterns in a more diverse range of nematode species, revealing significant variation in dye-filling between species and even between life stages within the same species (Koga and Tada, 2000). The authors found that staining patterns in parasitic species were often diminished compared to their free-living counterparts. This observation held true regardless of preferred host or invasion strategy, for example amphid neurons of the soybean cyst nematode, *Heterodera glycines*, and the root knot nematode *Meloidogyne hapla*, did not stain. In addition, no dye-filling was observed in the amphid neurons of the infectious juvenile (IJ) stage of the entomopathogenic nematode *Heterorhabditis bacteriophora*, and only periodic staining of these neurons was detected in non-IJ stages. Our results are consistent with these findings and suggest that, although the architecture of sensory anatomy is generally considered conserved across the phylum, modifications to sensory neurons exist between species and that these alterations may be a result of stage- and niche-specific adaptations.

To our knowledge, this is the first report to demonstrate that filarial nematodes exhibit specific tactile behaviors in response to host-derived odorants and the first to demonstrate chemosensory responses to host-derived compounds in *B. malayi*. Of note, both 3-methyl-1-butanol and 1-nonanol, which are human skin and sweat odorants and are known mosquito attractants (Meijerink *et al*., 2001; Qiu *et al*., 2011; Verhulst *et al*., 2011; Mukabana *et al*., 2012; Mathew *et al*., 2013), repelled L3 stage parasites. In contrast, physiologic saline that mimics the internal environment of the mosquito *(Aedes* saline) and human host compounds (L-lactic acid and human serum) were attractive. Together, these results indicate chemosensation in this nematode that may be important in transmission of this parasite by providing both positive and negative stimuli to encourage host penetration. In contrast to S. *stercoralis*, which actively seeks out a suitable host, *B. malayi* L3s are transmitted to human hosts through the bite of an infected mosquito. When the mosquito takes a blood meal, these worms migrate out of the proboscis onto the skin and crawl into the wound track left by the blood-feeding mosquito. In contrast, S. *stercoralis* must actively seek out a suitable host, so these parasites are strongly attracted to 3-methyl-1-butanol, 2-methyl-1-butanol and 1-nonanol, all of which are compounds found in human sweat (Meijerink *et al*., 2001; Castelletto *et al*., 2014). Unlike *B. malayi* L3s, IJ stage S. *stercoralis* are soil-dwelling nematodes, and it is likely that these and other host-derived compounds direct this parasite to a suitable host, where *B. malayi* is delivered directly onto the host skin and has to rapidly enter before it desiccates and dies. The results presented here indicate that *B. malayi* either have no response to, or are repelled by compounds found in human sweat. This negative response may facilitate transmission of the parasite by acting as a stimulant to move away from the host surface and drive skin penetration while the attractive response to serum and L-lactic acid may act as a "beacon” to direct the parasite to the wound track left by the mosquito vector.

This study describes a platform to interrogate filarial responses to external stimuli and may be further adapted to investigate other sensory cues such as temperature and pH. Historically, platforms used to investigate nematode sensory responses and pathways have leaned heavily on agar plate-based formats. Such assays have many advantages; they are simple, cost-effective, generate reasonable throughput, and are well suited to motile free-living, entomopathogenic and plant-parasitic species. Translating such simple assays to focus on animal-parasitic nematodes is more challenging but our work here is significant because it reinforces the idea that plate-based assays can be leveraged to ask fundamental questions about sensory perception in nematodes of veterinary and medical importance in a very straight forward manner. This plate-based assay is effective in identifying coarse tactic behaviors in *B. malayi* infective stage larvae - a stage that inherently is transitioning from the vector to the vertebrate host. The assay may not be well-suited for other life stages of the parasite, or for recognizing and quantifying more nuanced behaviors. As this work continues it may be possible to integrate other approaches to increase the precision and complexity of data generated. Recording electrophysiological activity from nematode sensory neurons is possible and a protocol has been described for immobilized individual adult *B. pahangi* (Perry, 2001; Rolfe *et al*., 2001). Such assays are technically demanding, however, and have limited throughput. Another exciting possibility is to adapt recent advances in microfluidic technologies that have been applied to *C. elegans* to create so-called "worm-on-a-chip” devices capable of handling and recording sensory responses in those nematodes (for review, see Bakhtina and Korvink, 2014). The design and fabrication of such devices is relatively simple and the low cost of manufacture combined with great opportunities for complexity, integration and functionality should allow these on-chip devices to be used with parasitic nematodes.

In summary, the research presented herein introduces a platform to better understand those molecular pathways underpinning filarial worm tactic behaviors central to transmission and infection, and positions investigators to define the precise role that chemosensation plays in these processes. Additional research will be needed to characterize the particular molecular mechanisms at play, and the assay we describe can be interfaced with reverse genetic approaches to provide a genetic and mechanistic basis for the observed responses. To this end, we have developed a robust and reliable in vivo RNAi protocol to suppress genes of interest in the *B. malayi* L3 stage (Song *et al*., 2010), whilst others have shown that RNAi can be applied to other life stages of the parasite in vitro (Aboobaker and Blaxter, 2003; Ford *et al*., 2009; Landmann *et al*., 2012; Singh *et al*., 2012; Winter *et al*., 2013; Luck *et al*., 2016; Misra *et al*., 2017; Verma *et al*., 2017). This receptiveness to RNAi, combined now with assays to interrogate sensory phenotypes, advances *B. malayi* as a tractable system to better understand sensory biology in filarial worm parasites. This may lead to independent or synergistic strategies to help control diseases of medical and veterinary importance.

## Acknowledgements

The authors would like to thank Brendan Dunphy, Michael Nazarchyk and Joel Bauer for technical assistance with mosquito rearing and maintenance. Parasite materials were provided by Dr. Andrew Moorhead, Erica Burkman, Molly Riggs and the NIH-NIAID Filariasis Research Reagent Resource Center (FR3) (www.filariasiscenter.org).

